# VMHdm/c^SF-1^ Neuronal Circuits Regulate Skeletal Muscle PGC1-α via the Sympathoadrenal Drive

**DOI:** 10.1101/2022.04.01.486756

**Authors:** Takuya Yoshida, Mina Fujitani, Scotlynn Farmer, Ami Harada, Zhen Shi, Jenny J. Lee, Arely Tinajero, Ashish K. Singha, Teppei Fujikawa

**Affiliations:** Department of Cellular and Integrative Physiology, Long School of Medicine, University of Texas Health San Antonio, San Antonio, US; Department of Clinical Nutrition School of Food and Nutritional Sciences, University of Shizuoka, Shizuoka, Japan; Center for Hypothalamic Research, Department of Internal Medicine, UT Southwestern Medical Center, Dallas, US; Laboratory of Nutrition Science, Department of Bioscience, Graduate School of Agriculture, Ehime University, Matsuyama, Japan; Nara Medical University, Nara, Japan; Department of Plastic Surgery, Hospital Zhejiang University School of Medicine, Zhejiang, China

## Abstract

**Objective:** To adapt to metabolically challenging environments, the central nervous system (CNS) orchestrates metabolism of peripheral organs including skeletal muscle. The organ-communication between the CNS and skeletal muscle has been investigated, yet our understanding of the neuronal pathway from the CNS to skeletal muscle is still limited. Neurons in the dorsomedial and central parts of the ventromedial hypothalamic nucleus (VMHdm/c) expressing steroidogenic factor-1 (VMHdm/c^SF-1^ neurons) are key for metabolic adaptations to exercise, including increased basal metabolic rate and skeletal muscle mass in mice. However, the mechanisms by which VMHdm/c^SF-1^ neurons regulate skeletal muscle function remain unclear. Here, we show that VMHdm/c^SF-1^ neurons increase the sympathoadrenal activity and regulate skeletal muscle peroxisome proliferator-activated receptor gamma coactivator 1 alpha (PGC-1α) in mice via multiple downstream nodes.

**Methods:** Optogenetics was used to specifically manipulate VMHdm/c^SF-1^ neurons combined with genetically-engineered mice and surgical manipulation of the sympathoadrenal activity.

**Results:** Optogenetic activation of VMHdm/c^SF-1^ neurons dramatically elevates mRNA levels of skeletal muscle *Pgc-1α*, which regulates a spectrum of skeletal muscle function including protein synthesis and metabolism. Mechanistically, the sympathoadrenal drive coupled with β2 adrenergic receptor (β2AdR) is essential for VMHdm/c^SF-1^ neurons-mediated increases in skeletal muscle PGC1-α. Specifically, both adrenalectomy and β2AdR knockout block augmented skeletal muscle PGC1-α by VMHdm/c^SF-1^ neuronal activation. Optogenetic functional mapping reveals that downstream nodes of VMHdm/c^SF-1^ neurons are functionally redundant to increase circulating epinephrine and skeletal muscle PGC1-α.

**Conclusions:** Collectively, we propose that VMHdm/c^SF-1^ neurons-skeletal muscle pathway, VMHdm/c^SF-1^ neurons→multiple downstream nodes→the adrenal gland→skeletal muscle β2AdR, underlies augmented skeletal muscle function for metabolic adaptations.

## 1. Introduction

The central nervous system (CNS) orchestrates the whole-body metabolism[1; 2]. Within the CNS, the hypothalamus plays a dominant role in the regulation of metabolic homeostasis in response to dynamic challenges such as hypoglycemia[3], cold-exposure[4], and exercise[5]. Our previous work articulates that neurons in the dorsomedial and central parts of ventromedial hypothalamic nucleus (VMHdm/c neurons) substantially contribute to metabolic adaptations to exercise training including augmented skeletal muscle mass and basal metabolic rate in mice[6]. Knockdown of steroidogenic factor-1 (SF-1)[7; 8] in VMHdm/c neurons hampers exercise-induced mRNA expression of *peroxisome proliferator-activated receptor gamma coactivator 1 alpha* (*Pgc-1α*) in skeletal muscle[6]. PGC-1α is a key transcriptional regulator that controls a broad range of genes related to glucose and fat metabolism, mitochondrial function, angiogenesis, and protein synthesis[9; 10]. Loss- or gain-of-function of PGC-1α in skeletal muscle dramatically changes skeletal muscle physiology as well as whole-body metabolism[11; 12]. These data suggest that VMHdm/c neurons expressing SF-1 (VMHdm/c^SF-1^ neurons) mediate metabolic responses of skeletal muscle to exercise, thereby contributing to metabolic benefits of exercise. However, the mechanisms by which VMHdm/c^SF-1^ neurons mediate exercise-induced augmented skeletal muscle PGC-1α expression remains unclear. In particular, the pathway from VMHdm/c^SF-1^ neurons to skeletal muscle has yet to be unraveled.

PGC-1α expression in skeletal muscle is augmented by a variety of physiological stimuli[9; 13]. For example, *ex vivo* muscle contraction is sufficient to increase mRNA levels of *Pgc-1α* by activation of calcium signaling pathways[14; 15]. Notably, the adrenergic activation such as epinephrine, norepinephrine, and β-2 adrenergic receptors (β2AdR) agonist can dramatically increase mRNA levels of *Pgc-1α* in skeletal muscle[16; 17]. In contrast, blocking the adrenergic signaling by a systemic injection of β2AdR antagonist significantly hampers exercise-induced *Pgc-1α* mRNA in skeletal muscle[18]. Numerous studies have indicated that VMH neurons affect the sympathetic nervous system (SNS) activity[19; 20]. Knockdown of SF-1 in VMHdm/c neurons suppresses exercise-induced epinephrine release[6]. These data indicate that VMHdm/c^SF-1^ neurons regulate skeletal muscle PGC-1α via the SNS. However, the functional neurocircuits underlying VMHdm/c^SF-1^ neuronal regulation of the SNS are still unclear. A genetic labeling study using *Sf-1*-Cre mice, which express Cre-recombinase in VMHdm/c in adults, portraits that VMHdm/c neurons highly innervate to several areas that regulate the SNS activity, including the anterior bed nucleus of the stria terminalis (aBNST), preoptic area (POA), anterior hypothalamus area (AH), paraventricular hypothalamic nucleus (PVH), and periaqueductal gray (PAG)[21]. Although studies using optogenetics have identified that distinct downstream sites of VMHdm/c^SF-1^ neurons regulate blood glucose[22], defensive behaviors[23], and food intake[24], the vital downstream node of VMHdm/c^SF-1^ neurons regulating the SNS is unknown.

In this study, we used optogenetics and genetically-engineered mice to determine the key downstream nodes of VMHdm/c^SF-1^ neurons that regulate skeletal muscle PGC-1α via the SNS. We found that epinephrine release by the sympathoadrenal activity coupled with β2AdR is essential for VMHdm/c^SF-1^ neuronal-induced skeletal muscle PGC-1α. Furthermore, our results demonstrated that VMHdm/c^SF-1^ neurons regulate the SNS through functionally redundant circuits with the PVH and PAG acting as the main downstream nodes.

## 2. Material and Methods

### 2.1 Genetically-Engineered Mice

*Sf-1*-Cre mice were obtained from the Jackson Laboratory (US; Strain# 012462, RRID:IMSR_JAX:012462). *Abrb2* KO mice (RRID:IMSR_JAX:031496) were derived from β-null mice[25], which was kindly provided by Dr. Bradford Lowell (Harvard Medical School). The sequences of genotyping primers are followings; for *Sf-1*-Cre, aggaagcagccctggaac, aggcaaattttggtgtacgg, agaaactgctccgctttcc with expected bands sizes of 627 bp for internal control and 236 bp for *Sf-1*-Cre; for *Abrb2* knockout (KO), cacgagactagtgagacgtg, accaagaataaggcccgagt, ccgggaatagacaaagacca with expected bands sizes of 225 bp for the wild-type allele and 410 bp for the knockout allele*. Sf-1*-Cre and *Abrb2* KO mice were bred to generate *Sf-1*-Cre::*Abrb2*^KO/KO^ and *Sf-1*-Cre::*Abrb2*^WT/WT^ mice. *Sf-1*-Cre mice were on a C57BL/J6 background and other mice are on a mixed background (C57BL/J6 and FVB.129). We used 3-6 month-old male mice whose body weights were above approximately 25-30 grams. Mice were housed at room temperature (22– 24 °C) with a 12 hr light/dark cycle (lights on at 6am, and 7am during daylight saving time) and fed a normal chow diet (2016 Teklad global 16% protein rodent diets, Envigo, US). Mice were maintained in groups and singly housed after adeno-associated virus (AAV) injections and optic fiber probes insertions. Animal care was according to established NIH guidelines, and all procedures were approved by the Institutional Animal Care and Use Committee of the University of Texas Southwestern Medical Center and University of Texas Health San Antonio.

### 2.2 AAV injections and optic fiber probe insertions

Recombinant AAVs were purchased from the Vector Core at the University of North Carolina at Chapel Hill, US or Addgene Inc (US). rAAV5-EF1α-DIO-mCherry (3.3 x 10^12^ VM/mL), rAAV5-EF1α-DIO-channelrhodopsin2(ChR2)(H134R)-mCherry (3.4 x 10^12^ VM/mL), and rAAV5-EF1α-DIO-ChR2(H134R)-EYFP (3.2 x 10^12^ VM/mL) were unilaterally administered into the right side of the VMH of mice using a UMP3 UltraMicroPump (WPI, US) with 10 µL NanoFil microsyringe (WPI) and 35G NanoFil beveled needle (WPI; Catalog# NF35BV-2). The volume of AAVs was 500-1000 nL at the rate 50-100 nL per minute, and the needle was left for another 5 minutes after the injection was finished. The face of beveled needle was placed towards the center of the brain. The coordinates of VMH-microinjection were AP; −1.4 L +0.5, and D −5.5 (from Bregma). The optic fiber probe was inserted as following coordinates; the VMH (AP; −1.4, L +0.5, and D −5.0), aBNST (AP; 0.3, L +0.5, and D −3.75), POA (AP; 0.4, L +0.25, and D −4.5), AH (AP; −0.9, L +0.5, and D −4.75), PVH/AH (AP; −0.8, L +0.25, and D −4.5), and PAG (AP; −4.3, L +0.2, and D −2.0). The configuration of the probe for the VMH stimulation was 400 µm Core, 0.5NA, Ø2.5 mm ceramic ferrule, and 6mm length (RWD Life science Inc, US and Doric Lenses, Canada) for the LED input. The configuration of probe for other sites and the VMH (S Figure 4) was 200 µm Core diameter, 0.39NA, Ø2.5 mm Ceramic Ferrule diameter, and with varied length varied (3-6 mm depending on the place) (RWD Life science Inc, US and Doric Lenses, Canada). The fiber probe was secured by adhesion bond (Loctite 454, Loctite Inc, US). Mice were allowed to recover for at least three weeks after the AAV injections to fully express recombinant proteins.

### 2.3. Optogenetics

The light emitting diode (LED) driver (Thorlabs, US; DC4104) with fiber-coupled LED (Thorlabs; Catalog# M470F3) was used for the VMH stimulation, and the laser unit (Opto engine LLC, US; Catalog# MDL-III-470) was used for the terminal of VMHdm/c^SF-1^ neuronal stimulations. The power of tips was set to ∼1 mW/mm^2^ for the VMH stimulation and ∼10 mW/mm^2^ for the terminal stimulations. The customed transistor-transistor logic generator was built based on the design by the University of Colorado Optogenetics and Neural Engineering Core. Rotary joint patch cable (Thorlabs; Catalog# RJPSF4 for LED and RJPFF4 for the laser) was used to connect either fiber-coupled LED or the laser unit. The quick-release interconnect (Thorlabs; Catalog# ADAF2) was used to connect the rotary joint patch cable to the fiber probe attached to mouse head. The stimulation was set to; 5 ms duration, 20 Hz, 2 seconds activation and 2 seconds rest cycle, 30 minutes (Figure 1B) otherwise specified.

**Figure 1.**
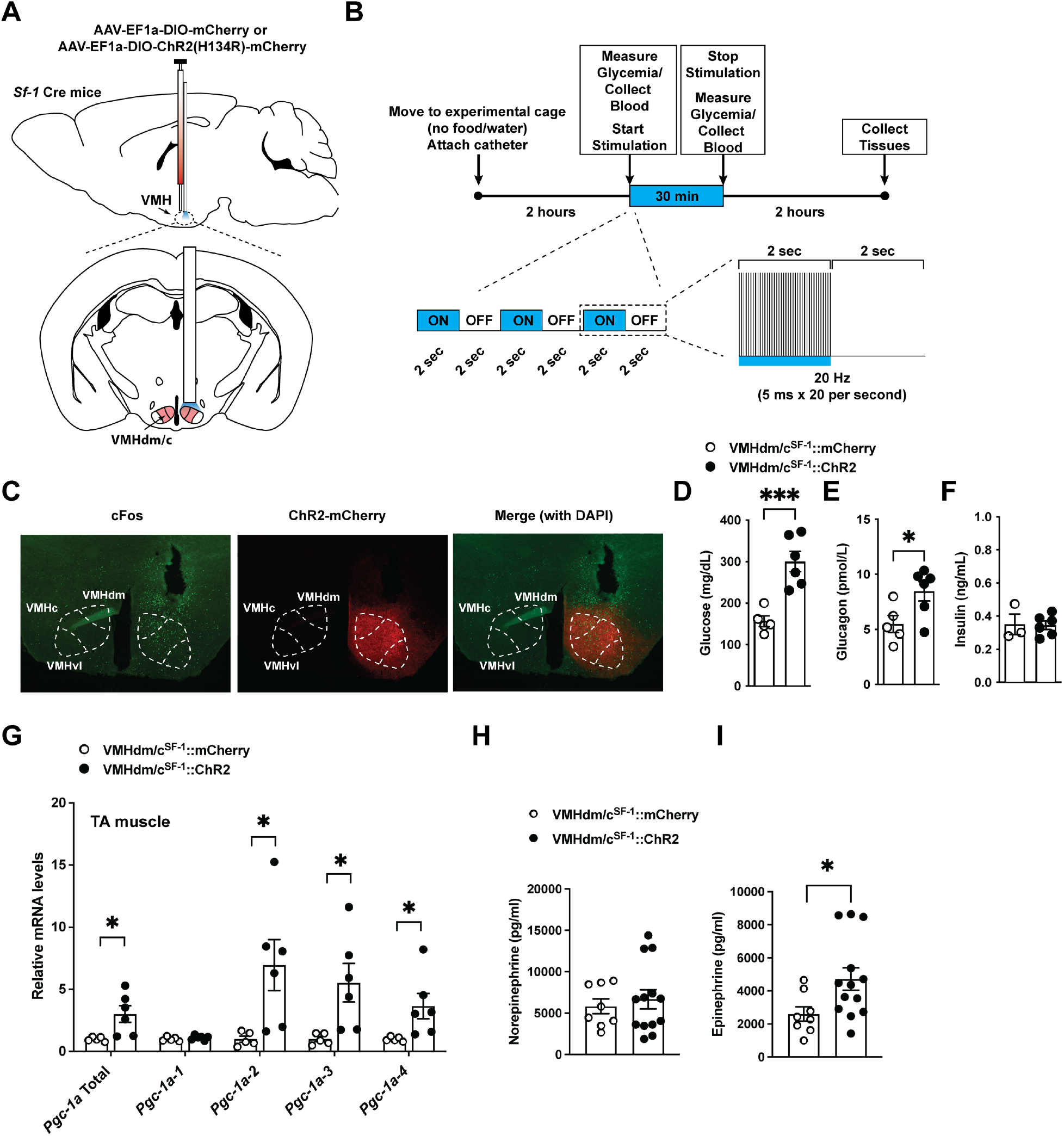
Optogenetic activation of VMHdm/c^SF-1^ neurons increases mRNA levels of skeletal muscle *Pgc1-α*. (**A**) Schematic figure of targeting site of VMHdm/c^SF-1^ neurons. (**B**) Experimental design and stimulus setting of optogenetics. (**C**) Representative figures of Fos expression pattern in the hypothalamus of mice expressing ChR2 in SF-1 expression neurons in VMHdm/c (VMHdm/c^SF-1^::ChR2) after VMHdm/c^SF-1^ neurons stimulation. (**D**) Blood glucose, (**E**) plasma glucagon, (**F**) plasma insulin in mice after VMHdm/c^SF-1^ neuronal stimulation. Control group is composed of mice expressing mCherry in VMHdm/c^SF-1^ neurons (VMHdm/c^SF-1^::mCherry) (**G**) mRNA expression levels of *Pgc1-α* isoform in skeletal muscle of VMHdm/c^SF-1^::ChR2 mice after optogenetic stimulation. (**H**) Plasma norepinephrine and (**I**) epinephrine in VMHdm/c^SF-1^::ChR2 mice after optogenetic stimulation. Values are mean ± S.E.M. *** p < 0.001, * p < 0.05.

### 2.4 Chemogenetics

Gi-Designer Receptors Exclusively Activated by Designer Drugs (DREADDs)-AAV (Gi-DREADD), rAAV8-hSyn-DIO-hM4D(Gi)-mCherry (Addgene Inc, Catalog# 44362-AAV8, 3.2 x 10^12^ VM/mL), was used for inhibition of VMHdm/c^SF-1^ neuronal activity induced by optogenetics. rAAV8-hSyn-DIO-hM4D(Gi)-mCherry and rAAV5-EF1α-DIO-ChR2(H134R)-EYFP (250 nL each, total 500 nL) were injected into the right side of the VMH of *Sf-1*-Cre mice as described above. Control for rAAV5-EF1α-DIO-ChR2(H134R)-EYFP was rAAV5-EF1α-DIO-EYFP. The optic cannula was inserted the VMH, PVH/AH or PAG as described above. Clozapine-N-Oxide (CNO, catlog# HB6149, Hello Bio Inc, NJ, USA) was injected intraperitoneally (i.p.) 10 minutes before optogenetic stimulation begun (2.5 mg/kg per BW). Sterile saline was used as vehicle and control.

### 2.5 Adrenalectomy (ADX)

Skin and muscle incisions (approximately 1 cm) were made close to the abdominal area where kidney was located. Both the adrenal glands were removed by tying with sterile suture. The sham operation was performed as same as ADX surgery except for removing the adrenal glands. The corticosterone water (75 µg/mL, vehicle was sterile 1% ethanol/0.9% NaCl water) was provided with ADX mice, and vehicle was provided with sham mice.

### 2.6 Immunohistochemistry and Fos counting

Mouse brains were prepared as previously described[26; 27]. Anti-cFos (Sigma, US; Catalog# F7799, batch #0000102540, RRID:AB_259739) and secondary fluorescent antibodies (Thermofisher Inc, US; Catalog# A-21202, Lot#WF320931, and RRID:AB_141607; A-21203, Lot#WD319534, and RRID:AB_141633; A10523, Lot#, and RRID:AB_1500679) were used. Dilution rates for antibodies were 1:1000 for first antibody and 1:200 for second antibodies. Images were captured by fluorescence microscopies (Keyence US, US; Model: BZ-X710, Leica Inc, US; Model DM6 B). Exposure of captured images was adjusted, and each area (aBNST, POA, AH, PVH, VMH, and PAG) was clipped by Photoshop based on the mouse brain atlas[28]. Clipped images were exported to Fiji, and the number of cells expressing Fos was counted by particle measurement function (Figure 5B).

### 2.7 Assessment of glucose, catecholamines, and hormone levels in the blood

Blood glucose was measured by a glucose meter as previously described[6; 27; 29]. Plasma catecholamines and hormones levels were measured as previously described[6; 27]. Briefly, the plasma catecholamines were analyzed by the Vanderbilt Hormone Assay & Analytical Services Core. Plasma Glucagon (Mercodia Inc, US; Catalog#10-1281-01), insulin (Crystal Chem Inc, US; Catalog# 90080), and corticosterone (Cayman Chemical, US; Catalog #501320) levels were determined by commercially available ELISA kits.

### 2.8 Assessment of mRNA

mRNA levels in the tibialis anterior (TA) muscle were determined as previously described[30]. The sequences of the deoxy-oligonucleotides primers are: *Ppargc1a* total (5’ tgatgtgaatgacttggatacagaca, and 5’ gctcattgttgtactggttggatatg), *Ppargc1a*-*1* (5’ ggacatgtgcagccaagactct, and 5’ cacttcaatccacccagaaagct), *Ppargc1a*-*2* (5’ ccaccagaatgagtgacatgga, and 5’ gttcagcaagatctgggcaaa), *Ppargc1a*-*3* (5’ aagtgagtaaccggaggcattc, and 5’ ttcaggaagatctgggcaaaga), *Ppargc1a*-*4* (5’ tcacaccaaacccacagaaa, and 5’ ctggaagatatggcacat), and *18S* (5’ catgcagaacccacgacagta and 5’ cctcacgcagcttgttgtcta).

### 2.9 Data analysis and statistical design

The data are represented as means ± S.E.M. GraphPad PRISM version 9 (GraphPad, US) was used for the statistical analyses and *P*<0.05 was considered as a statistically significant difference. A detailed analysis was described in Supplemental Table 1. The sample size was decided based on previous publications[6; 26; 27; 29-34], and no power analysis was used. Experiments were biologically replicated in Figure 1, 2, 3, and Figure 4B. Experiments for qPCR were technically replicated. We did not carry out replicate experiments for data shown in Figure 4C-H and 5. Figures were generated by PRISM version 9, Illustrator 2021-23, and Photoshop 2021-23 (Adobe Inc, US).

## 3. Results

### 3.1 Optogenetic VMHdm/c^SF-1^ neuronal activation induces skeletal muscle *Pgc-1α* mRNA expression

To determine the neuronal mechanism by which VMHdm/c^SF-1^ neurons regulate skeletal muscle function via the SNS, we generated mice expressing ChR2(H134R)[35] specifically in VMHdm/c^SF-1^ neurons by microinjection of AAV bearing Cre-dependent ChR2 fused with fluorescent reporters (AAV-DIO-ChR2) into *Sf-1* Cre mice[36] (VMHdm/c^SF-1^::ChR2, Figure 1A). We used AAV containing Cre-dependent mCherry for control (VMHdm/c^SF-1^::mCherry). We used the following stimulation configurations; 5 ms duration, 20 Hz, 2 seconds activation/2 seconds resting cycle for 30 minutes (Figure 1B). We confirmed that our optogenetic configuration significantly induced Fos protein expression, a neuronal activation marker, in the stimulated side of the VMHdm/c, but not in the non-stimulated side (Figure 1C). Similar to previous studies[22; 37; 38], VMHdm/c^SF-1^ neuronal activation increased blood glucose (Figure 1D). In addition, we observed that VMHdm/c^SF-1^ neuronal activation induced increases in plasma glucagon without changing plasma insulin levels (Figure 1E and F). Next, we determined whether VMHdm/c^SF-1^ neuronal activation can induce transcriptional changes in skeletal muscle. PGC-1α is a key transcriptional regulator for a spectrum of genes governing glucose and fat metabolism, oxidative capacity, protein synthesis and degradation, and vascularization[9; 10; 13]. Importantly, *Pgc-1α* mRNA induction can be used as a readout of exercise-related skeletal muscle transcriptional changes because a single exercise training dramatically increases mRNA levels of *Pgc-1α[6; 13]*. PGC-1α has several isoforms such as PGC1a-1, -2, -3 and -4[39]. We found that 30 minutes of VMHdm/c^SF-1^ neuronal activation is sufficient to induce *Pgc-1α-2, -3,* and *-4* mRNA levels in TA skeletal muscle (Figure 1G). We then measured plasma catecholamines levels after VMHdm/c^SF-1^ neuronal activation. Interestingly, plasma epinephrine levels were significantly increased by VMHdm/c^SF-1^ neuronal activation, but norepinephrine levels in VMHdm/c^SF-1^ neuronal activated mice were not significantly different from control group (Figure 1 H and I). The increased epinephrine likely caused suppression of insulin secretion while mice showed hyperglycemia. Previous studies have shown that optogenetic VMHdm/c^SF-1^ neuronal activation elicits behavioral changes such as freeze and burst activity (combination of freeze, jump, and run)[23; 40]. In line with that, we found that VMHdm/c^SF-1^ neuronal activation induced freeze or burst activity (17 cases of freeze (55%), 14 cases of burst activity (45%); total 31 trials). However, we did not observe any differences of skeletal muscle *Pgc-1α* mRNA levels between mice showed freeze and burst activity (SFigure 1), suggesting that the burst activity is unlikely to contribute to VMHdm/c^SF-1^ neurons-induced *Pgc-1α* expression. Collectively, these data indicate that VMHdm/c^SF-1^ neurons regulate skeletal muscle function via sympathoadrenal activity, as the adrenal gland is the only organ able to secret epinephrine into circulation.

### 3.2 The adrenal gland is essential for VMHdm/c^SF-1^ neuronal regulation of skeletal muscle

Next, to determine whether epinephrine release from the adrenal gland is required for VMHdm/c^SF-1^ neurons-induced skeletal muscle *Pgc-1α*, we surgically removed the adrenal gland, and then stimulated VMHdm/c^SF-1^ neurons. Sham surgery were used as surgical controls. As we expected, the ADX eliminated blood corticosterone (Figure 2B). ADX mice were supplied with corticosterone in the drinking water to maintain the physiological levels of blood corticosterone (Figure 2C). We carried out optogenetic stimulation five days after ADX. ADX appeared not to affect Fos induction in the VMH (Figure 2D), suggesting ADX does not affect the ability of VMHdm/c^SF-1^ neuronal activation. Indeed, plasma glucagon levels were significantly higher in ADX mice after VMHdm/c^SF-1^ neuronal activation (Figure 2E). The removal of the adrenal medulla in rats blocks hyperglycemia induced by microinjection of a cholinergic agonist into the VMH[41]. In line with the report, VMHdm/c^SF-1^ neuronal activation could not increase blood glucose levels in ADX mice at all (Figure 2F). Stunningly, ADX also completely blocked the effects of VMHdm/c^SF-1^ neuronal activation on skeletal muscle *Pgc-1α* expression (Figure 2G), suggesting that the sympathoadrenal drive, specifically the epinephrine release, is essential for VMHdm/c^SF-1^ neuronal regulation of skeletal muscle function. In addition, these data suggest that glucagon release unlikely contributes to VMHdm/c^SF-1^ neuronal-induced blood glucose levels.

**Figure 2.**
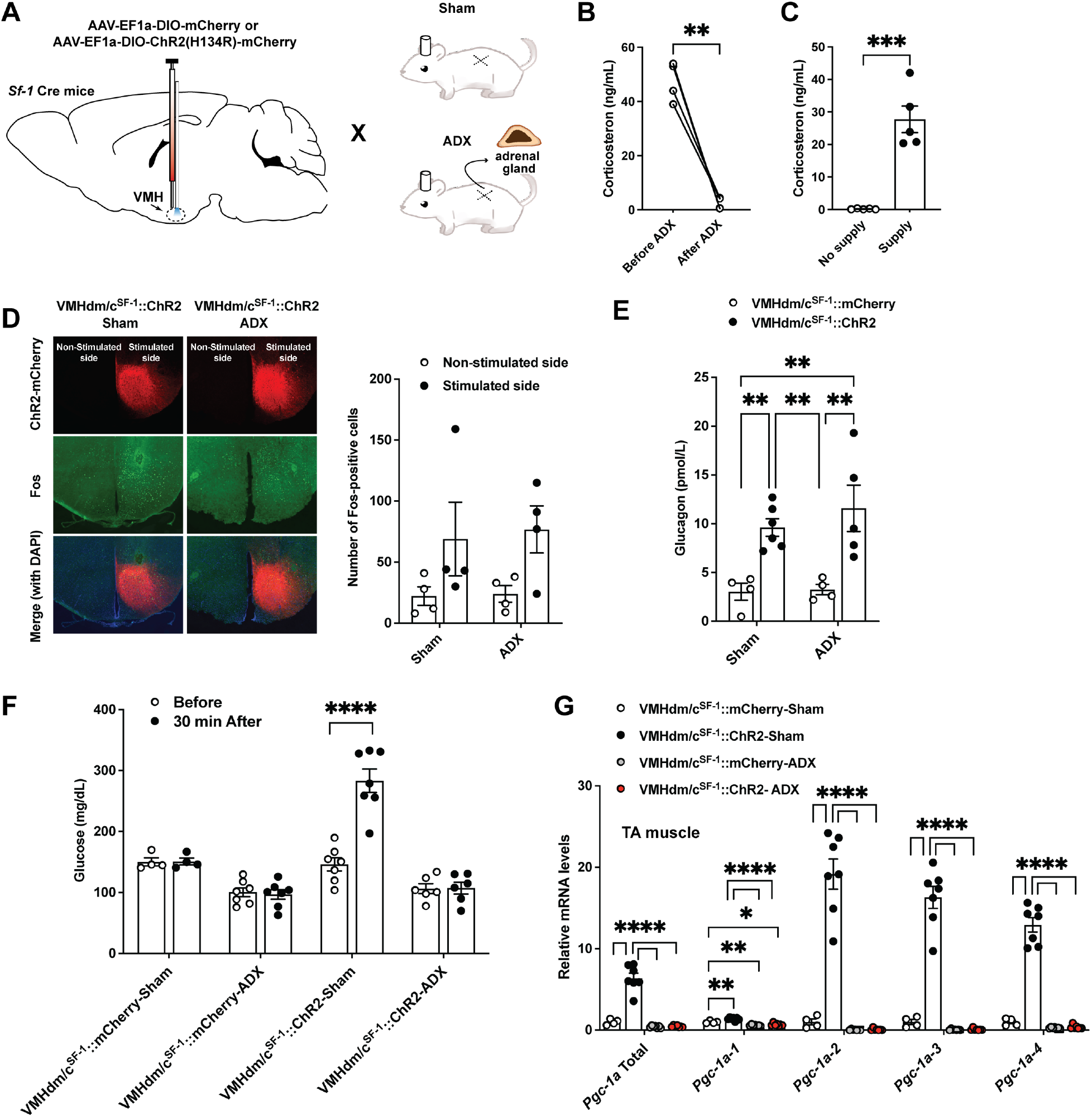
Adrenalectomy (ADX) completely blocks VMHdm/c^SF-1^ neurons-induced blood glucose and skeletal muscle *Pgc1-α* expression. (**A**) Schematic of ADX experimental design. (**B**) Blood corticosterone levels before and after ADX in mice. (**C**) Blood corticosterone levels in ADX mice with and without corticosterone supplementation. (**D**) Representative figures of Fos expression pattern in the hypothalamus of ADX mice expressing ChR2 in SF-1 expression neurons in VMHdm/c (VMHdm/c^SF-1^::ChR2-ADX) and the number of Fos positive cells in the VMH of VMHdm/c^SF-1^::ChR2-ADX and sham VMHdm/c^SF-1^::ChR2 (VMHdm/c^SF-1^::ChR2-sham). (**E**) Blood glucose, (**F**) plasma glucagon, and (**G**) mRNA expression levels of *Pgc1-α* isoform in skeletal muscle of VMHdm/c^SF-1^::ChR2-ADX after optogenetic stimulation. Values are mean ± S.E.M. **** p < 0.0001, *** p < 0.001, ** p <0.01, * p < 0.05.

### 3.3 β2AdR is required for VMHdm/c^SF-1^ neuronal-induced skeletal muscle *Pgc-1α*

We assessed the contribution of β2AdR in VMHdm/c^SF-1^ neuronal-induced skeletal muscle *Pgc-1α* expression. β2AdR is the major form of adrenergic receptors in skeletal muscle[42]. β2AdR agonist injection increases skeletal muscle *Pgc-1α-2, -3,* and *-4* mRNA levels[16], suggesting that the sympathetic input coupled with β2AdR is key for skeletal muscle physiology. We tested whether β2AdR global KO can diminish VMHdm/c^SF-1^ neuronal-induced skeletal muscle *Pgc-1α* expression (Figure 3A). We first determined whether β2AdR KO affects the ability of ChR2 to activate VMHdm/c^SF-1^ neurons. Optogenetic stimulation induced equal levels of Fos expression in the VMH of wild-type and β2AdR KO mice (VMHdm/c^SF-1^::ChR2-WT and VMHdm/c^SF-1^::ChR2-β2AdR^KO^, Figure 3B), suggesting β2AdR KO does not affect the capacity for optogenetic activation of VMHdm/c^SF-1^ neurons. Blood glucose and plasma glucagon in VMHdm/c^SF-1^::ChR2-β2AdR^KO^ were significantly higher than that of control group (VMHdm/c^SF-1^::mCherry-β2AdR^KO^) (Figure 3C and D). In addition, optogenetic stimulation of VMHdm/c^SF-1^ neurons significantly increased plasma epinephrine in β2AdR KO compared to non-stimulation control (Figure 3E). VMHdm/c^SF-1^::ChR2-β2AdR^KO^ showed significantly lower mRNA levels of skeletal muscle *Pgc-1α-2, -3,* and *-4* compared to VMHdm/c^SF-1^::ChR2-WT (Figure 3F). Collectively, these results demonstrate that β2AdR is essential for VMHdm/c^SF-1^ neuronal-induced skeletal muscle transcriptional regulation, but not blood glucose levels.

**Figure 3.**
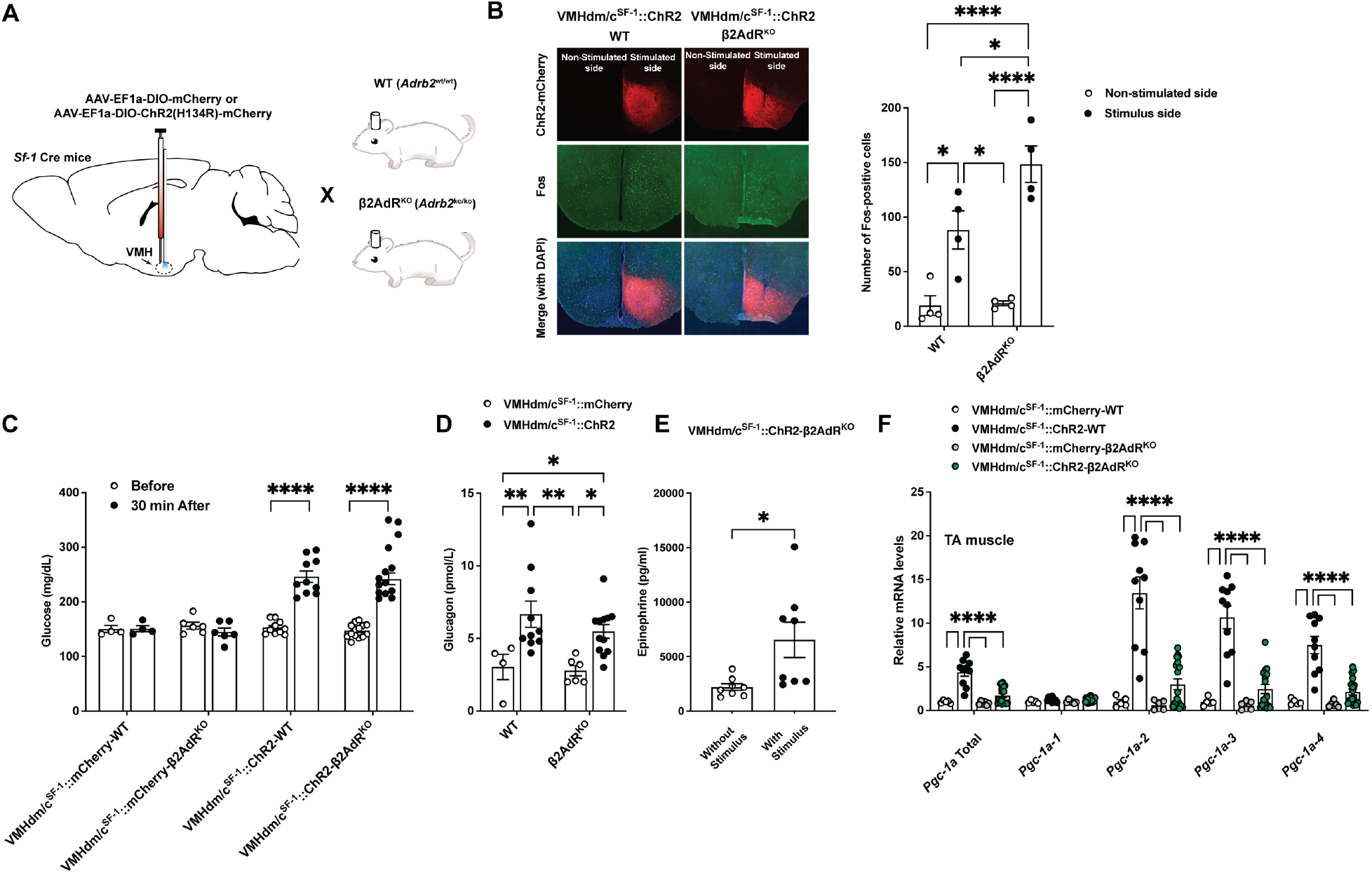
Ablation of β2AdR hampers VMHdm/c^SF-1^ neurons-induced skeletal muscle *Pgc1-α*. (**A**) Schematic of experimental design. (**B**) Representative figures of Fos expression pattern in the hypothalamus of VMHdm/c^SF-1^::ChR2 mice lacking β2AdR (VMHdm/c^SF-1^::ChR2-β2AdR^KO^), and the number of Fos positive cells in the VMH of VMHdm/c^SF-1^::ChR2-β2AdR^KO^ and wild-type control (WT) (VMHdm/c^SF-1^::ChR2-WT). (**C**) Blood glucose and (**D**) plasma glucagon levels in VMHdm/c^SF-1^::ChR2-β2AdR^KO^ mice after optogenetic stimulation. (**E**) Plasma epinephrine levels in VMHdm/c^SF-1^::ChR2-β2AdR^KO^ mice with and without optogenetic stimulation. (**F**) mRNA expression levels of *Pgc1-α* isoform in skeletal muscle of ChR2-β2AdR^KO^ mice after optogenetic stimulation. Values are mean ± S.E.M. **** p < 0.0001, ** p <0.01, * p < 0.05.

### 3.4 Redundant functionality of downstream nodes of VMHdm/c^SF-1^ neurons

VMHdm/c^SF-1^ neurons project to a broad range of the CNS sites that regulate the SNS including presympathetic nodes[21]. Among these areas, we examined the contributions of the aBNST, POA, AH, PVH/AH, and PAG to VMHdm/c^SF-1^ neuronal-induced augmentations in plasma epinephrine, and skeletal muscle *Pgc-1α* expression. As the PVH and AH are located in close proximity to each other (the AH is located at ventral side of the PVH), thus the light source can reach to both areas based on the theoretical irradiance value calculation (https://web.stanford.edu/group/dlab/cgi-bin/graph/chart.php), it is possible that the PVH stimulation also affects the AH. Therefore, going forward, we will refer to optogenetic stimulation in this region as the PVH/AH stimulation instead of the PVH stimulation. We injected AAV-DIO-ChR2 into the VMH of *Sf-1* Cre mice followed by the insertion of the optic fiber probe into each area (Figure 4A). Intriguingly, all terminal stimulations of VMHdm/c^SF-1^ neurons significantly increased blood glucose (Figure 4B). While stimulation in each area also increased plasma epinephrine (mean values ± SEM; 1919 ± 324, 3869 ± 922, 3782 ± 594, 2759 ± 188, 5145 ± 841, and 6311 ± 836 pg/mL for mCherry, aBNST, POA, AH, PVH/AH, and PAG, respectively), the most striking differences were observed after terminal stimulation of VMHdm/c^SF-1^ neurons in the PVH/AH and PAG (Figure 4C). We found that *Pgc-1α* total mRNA expression levels were significantly increased in the terminal stimulation of the POA and PVH/AH (Figure 4D). *Pgc-1α-1* mRNA expression levels were significantly increased in the terminal stimulation in the AH and PVH/AH (Figure 4E). *Pgc-1α-2* and *-3* mRNA expression levels were significantly increased in the terminal stimulation in the PVH/AH and PAG (Figure 4G). Finally, *Pgc-1α-4* mRNA expression levels were significantly increased in the terminal stimulation in the PVH/AH. Of note, although the terminal stimulation in the aBNST, POA, and AH did not statistically increase RNA levels of skeletal muscle *Pgc-1α-2, -3,* and *-4*, the mean values were higher than those in the control group. Collectively, we conclude that VMHdm/c^SF-1^ neuronal projections to the PVH/AH and PAG have greater contributions to the regulation of skeletal muscle *Pgc-1α* mRNA expression and plasma epinephrine. However, other projection sites likely contribute as well, indicating that VMHdm/c^SF-1^ neurons use neurocircuits that are functionally redundant to regulate the sympathoadrenal activity.

**Figure 4.**
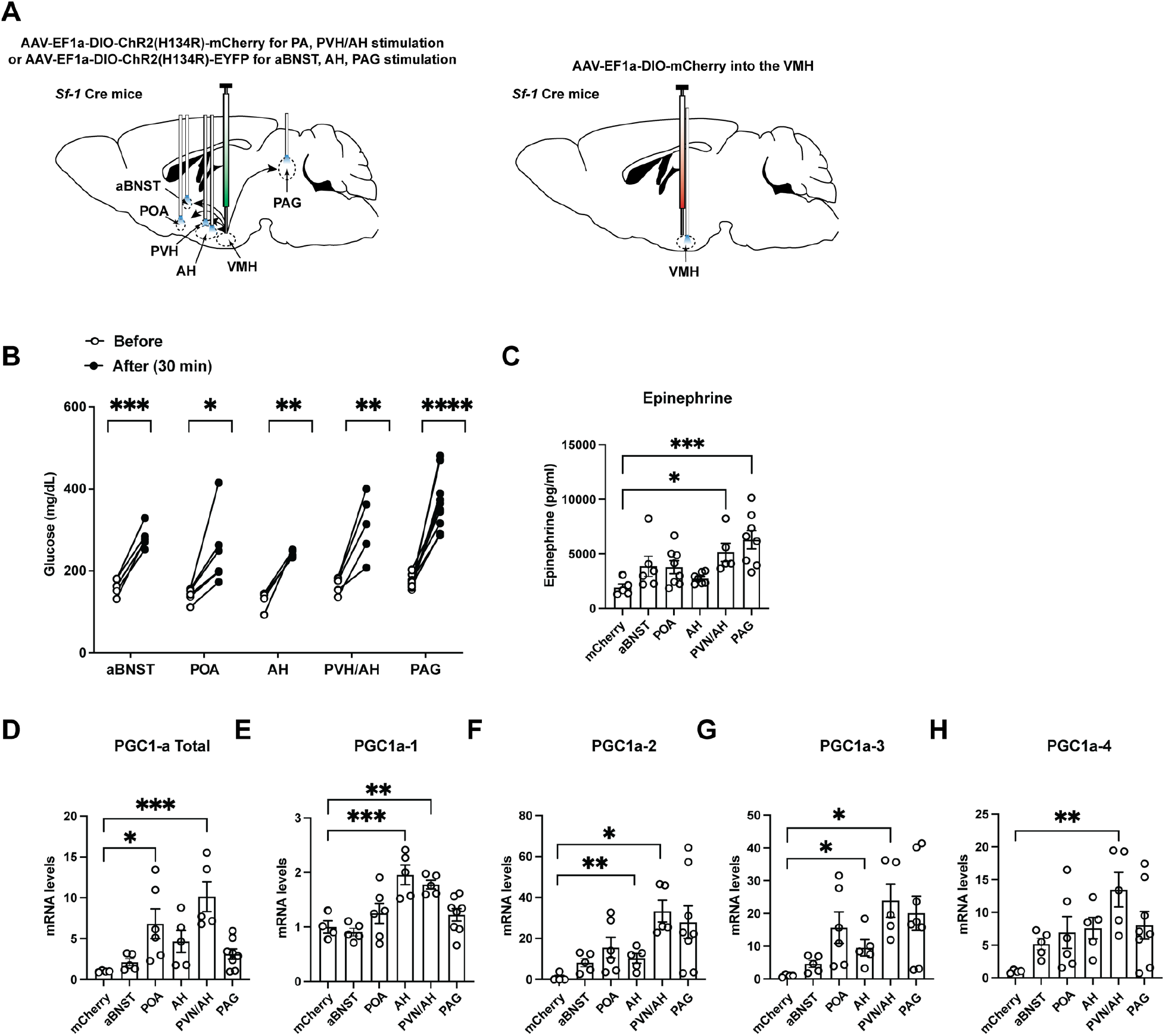
Downstream nodes of VMHdm/c^SF-1^ neurons are functionally redundant to regulate the sympathoadrenal activity. (**A**) Schematic figure of experimental design. (**B**) Blood glucose levels in VMHdm/c^SF-1^::ChR2 mice before and after the terminal stimulation of the anterior bed nucleus of the stria terminalis (aBNST), preoptic area (POA), anterior hypothalamus area (AH), paraventricular hypothalamic nucleus (PVH) and AH, and periaqueductal gray (PAG). Control group is VMHdm/c^SF-1^::mCherry mice after the VMH stimulation. (**C**) Plasma epinephrine levels in VMHdm/c^SF-1^::ChR2 mice after the terminal stimulation of aBNST, POA, AH, PVH/AH, and PAG. (**D**) mRNA expression levels of *Pgc1-α-Total*, (**E**) *Pgc1-α-1*, (**F**) *Pgc1-α-2*, (**G**) *Pgc1-α-3*, and (**H**) *Pgc1-α-4*, skeletal muscle of VMHdm/c^SF-1^::ChR2 mice after the terminal stimulation of aBNST, POA, AH, PVH/AH, and PAG. Values are mean ± S.E.M. **** p < 0.0001, *** p < 0.001, ** p <0.01, * p < 0.05.

A previous study found that VMHdm/c^SF-1^ neurons project to downstream sites collaterally[23]. For example, VMHdm/c^SF-1^ neurons projecting to the PAG also send axons to the AH[40]. Each VMHdm/c^SF-1^ neuron likely projects to multiple sites, especially to the areas that are topographically adjacent, such as aBNST and POA or the AH and PVH. In other words, when the terminal activation of VMHdm/c^SF-1^ neurons at one downstream area occurs, back-propagated neuronal activation could happen in other non-stimulated areas. To determine whether it is the case, we assessed Fos expression after the terminal activation of VMHdm/c^SF-1^ neurons in the aBNST, POA, AH, PVH/AH, and PAG as well as the soma stimulation of VMHdm/c^SF-1^ neurons (Figure 5A and B). We compared the stimulated side (right hemisphere) and the non-stimulated side (left hemisphere). As shown (Figure 1C), the optogenetic stimulation of the soma of VMHdm/c^SF-1^ neurons (Figure 5O and P, S Figure 2E) significantly increased Fos expression in the stimulated side of the VMH (Figure 5Q). Mice received the control AAV did not show any difference between stimulated and non-stimulated sides after the VMH stimulation (S Figure 3). Intriguingly, the soma activation of VMHdm/c^SF-1^ neurons significantly increased Fos expression only in the PVH and AH (Figure 5Q). We did not find distal neuronal propagations by the terminal activation in the POA, PVH/AH, AH, and PAG (Figure 5H, K, N, Q, and T, S Figure 2A, B, C, D and F). The terminal stimulation in the aBNST increased Fos expression in distal site PAG (Figure 5E). However, it is unclear that Fos expression in the PAG was induced by the distal propagation as Fos expression in the VMH was not changed after aBNST stimulation (Figure 5E). We frequently observed Fos-expression in the sites proximal to the stimulated site. For instance, the aBNST stimulation induced Fos-expression in the POA (Figure 5E). Likewise, the AH stimulation induced Fos-expression in the PVH (Figure 5K), and the PVH/AH stimulation induced Fos-expression in the POA and VMH (Figure 5N). Fos expression in the stimulated hemisphere of AH was significantly higher of all the stimulation sites (Figure 5E, H, K, N, Q, and T), suggesting that these Fos-inductions in the AH may be the secondary rather than direct propagated activation.

**Figure 5.**
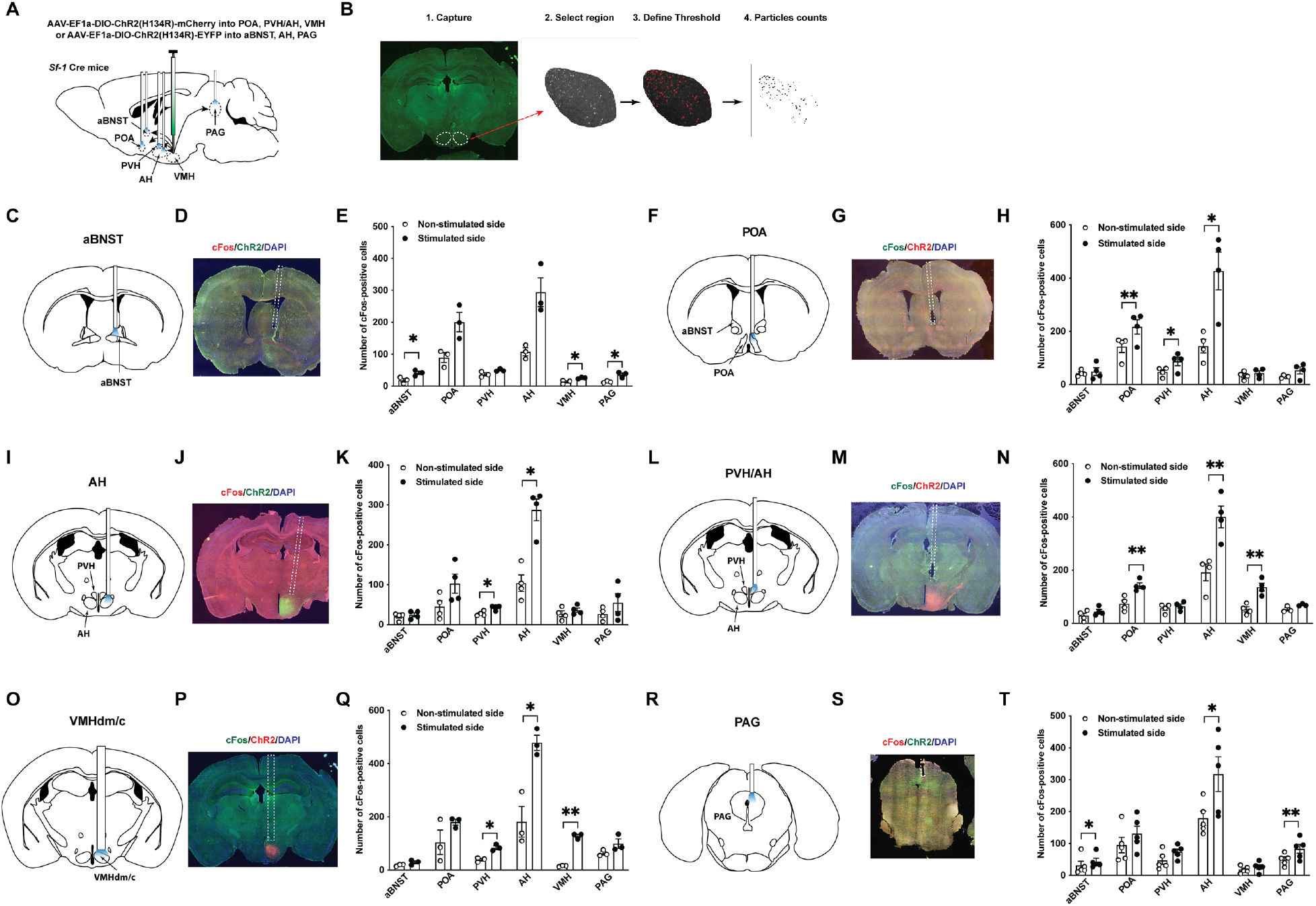
The terminal activation of VMHdm/c^SF-1^ neurons evokes Fos expression in the proximal terminal sites. (**A**) Schematic figure of experimental design. (**B**) Schematic of Fos expression counts in the nucleus. The brain maps of optic fiber insertion sites of (**C**), aBNST, (**F**) POA (**I**), AH (**L**), PVH/AH, (**O**), VMH, and (**R**) PAG. Representative figures of Fos expression at the terminal sites or soma of VMHdm/c^SF-1^ neurons; (**D**) aBNST, (**G**) POA, (**J**) AH, (**M**) PVH/AH, (**P**), VMHdm/c, and (**S**) PAG. The number of Fos expression cells at the aBNST, POA, AH, PVH/AH, and PAG of VMHdm/c^SF-1^::ChR2 mice after the terminal stimulation of (**E**) aBNST, (**H**) POA, (**K**) AH, (**N**) PVH/AH, (**Q**) VMHdm/c and (**T**) PAG. Values are mean ± S.E.M. **** p < 0.0001, *** p < 0.001, ** p <0.01, * p < 0.05.

To further investigate the redundant roles of downstream sites of VMHdm/c^SF-1^ neurons regulating the SNS, we stimulated the terminal of VMHdm/c^SF-1^ neurons in the PVH/AH or PAG while suppressing the soma of VMHdm/c^SF-1^ neurons to eliminate back-propagations, at least, between the PVH/AH and PAG. We selected these two sites as our data (Figure 4) showed that stimulation of those two sites led to the most prominent the SNS activation. We confirmed that Gi-DREADD hampers hyperglycemia, skeletal muscle *Pgc-1α* mRNA expression, and Fos activation induced by optogenetics (S Figure 4). The terminal stimulation of PVH/AH or PAG still induced hyperglycemia (S Figure 5A) and skeletal muscle *Pgc-1α* mRNA expression (S Figure 5B and C) while Gi-DREADD inhibited VMHdm/c^SF-1^ neurons. However, Gi-DREADD significantly suppressed *Pgc-1α* mRNA expression compared to control injection group (S Figure 5B and C). Gi-DREADD suppressed Fos activation in the VMH by the terminal stimulation of PVH/AH or PAG of VMHdm/c^SF-1^ neurons (S Figure 5D and F). As described above, we observed prominent Fos activation in the AH by any conditions, further confirming that these Fos expression in the AH may not be reflected to direct activation of VMHdm/c^SF-1^ neurons. The terminal stimulation of PVH/AH with Gi-DREADD did not induce Fos in the PAG, while the terminal stimulation of PAG with Gi-DREADD did not induce Fos in the VMH (S Figure 6 D and F). In these conditions, the readouts of the SNS activity (glucose and skeletal *Pgc-1α* mRNA expression) were still increased (S Figure 6A-C), suggesting that at least these two distal projection sites of VMHdm/c^SF-1^ neurons, PVH/AH and PAG projections, are functionally redundant. Collectively, Fos expression data indicated that each single VMHdm/c^SF-1^ neuronal axon projects many downstream sites collaterally, strengthening the idea of functionally redundant VMHdm/c^SF-1^ neuronal circuits that regulate the SNS.

## 4.1 Discussion

Here we demonstrate that VMHdm/c^SF-1^ neuronal circuits regulate skeletal muscle *Pgc-1α* mRNA via the sympathoadrenal-released epinephrine coupled with β2AdR. Our data further predict that VMHdm/c^SF-1^ neurons collaterally project multiple presympathetic nodes in the CNS that possess redundant functions to regulate the sympathoadrenal activation. The VMH facilitates glucose uptake in skeletal muscle and brown adipose tissue (BAT) and fatty acid mobilization from white adipose tissues (WAT) by direct sympathetic innervation rather than the sympathoadrenal drive[43–47]. Intriguingly, these studies have demonstrated that direct sympathetic innervation is key for VMH-induced glucose uptake and lipolysis in BAT and skeletal muscle rather than the sympathoadrenal activity[43; 44; 46; 47]. Contrary to these previous findings, results in this study highlight the importance of the sympathoadrenal activity for VMHdm/c^SF-1^ neurons-induced skeletal muscle *Pgc-1α* expression. Because VMH neurons are genetically heterogenous[48; 49], it is possible that VMHdm/c^SF-1^ neurons regulate skeletal muscle function primarily via sympathoadrenal activity, while other subtypes of neurons in the VMH mediate skeletal muscle glucose uptake via the direct sympathetic innervation. Optogenetic activation of leptin receptors-expressing neurons in the VMHdm/c does not affect blood glucose, plasma glucagon, and plasma insulin levels[22], supporting the notion that functional segregation may exist within the VMHdm/c neuronal subgroups. Further studies are warranted to reveal the mechanistic differences among the genetically distinctive neuronal groups within the VMHdm/c that regulate skeletal muscle function.

PGC-1α is one of the key molecules regulating a wide range of skeletal muscle physiology[50]. PGC-1α-1 is “classic” PGC-1α, and regulates mitochondrial function, glucose and fat metabolism[14; 39]. PGC-1α-4 regulates the protein synthesis[14; 39]. The physiological role of PGC-1α-2 and -3 are still not clear. It is predicted that they contribute to angiogenesis, epithelial function, chromosomal maintenance, and cholesterol metabolism[16]. PGC-1α-1 is derived from the proximal promoter region of PGC-1α locus, and PGC-1α-2, -3, and -4 are derived from the distal promoter region[10]. Intriguingly, running wheel activity and treadmill exercise activate the distal promoter region but not the proximal promoter region[18; 51]. Furthermore, β2AdR agonist also only initiates the transcription of PGC-1α exclusively at the proximal promoter region[18; 51], leading to increased mRNA levels of PGC-1α-2, -3, and -4, but not PGC-1α-1[16]. These data indicate the importance of the CNS→the SNS→skeletal muscle β2AdR axis for the regulation of skeletal muscle PGC-1α in response to exercise. Our data demonstrate that VMHdm/c^SF-1^ neurons can regulate PGC-1α-2, -3, and -4, but not -1, further strengthening the idea that these neurons are important for exercise-induced augmentation of skeletal muscle function.

VMHdm/c^SF-1^ neuronal activation increases circulating epinephrine, and ADX completely blocks VMHdm/c^SF-1^ neuronal-induced skeletal muscle *Pgc-1α* expression (Figure 2), suggesting that epinephrine secretion from the adrenal grand is essential for VMHdm/c^SF-1^ neuronal regulation of skeletal muscle function. Because the adrenal gland secretes many endocrine hormones, we must consider the contribution of non-catecholamines hormones to VMHdm/c^SF-1^ neuronal-induced skeletal muscle *Pgc-1α* expression. For instance, optogenetic activation of VMHdm/c^SF-1^ neurons increases blood corticosterone[22]. Thus, it is possible that the surge of corticosterone from the adrenal gland also contributes to VMHdm/c^SF-1^ neuronal-induced skeletal muscle *Pgc-1α*. Corticosterone supplement (Figure 2B) can only provide physiologically stable corticosterone levels but can not imitate endogenous dynamics of corticosterone releases. However, dexamethasone (DEX) treatments suppress PGC-1α in skeletal muscle cell lines[52], and in the testis of *in vivo* mouse model[53]. Glucocorticoid receptors (GRs) are expressed in skeletal muscle, and skeletal muscle-specific GR-KO mice show increased lean mass[54], which is in line with the fact that DEX can induce atrophy[55]. Therefore, we conclude that corticosterone unlikely contributes to the induction of skeletal muscle PGC-1α after VMHdm/c^SF-1^ neuronal activation. However, further studies are necessary to determine that epinephrine is the only factor in the adrenal gland that contributes to VMHdm/c^SF-1^ neuronal-induced skeletal muscle *Pgc-1α*.

A large body of literature has concluded that β2AdR signaling is key to the regulation of skeletal muscle PGC-1α expression[42], and ultimately orchestrating protein synthesis/degradation, glucose and fat metabolism, and angiogenesis[9; 10; 51]. Our study extends these findings to show that the sympathoadrenal activity coupled with β2AdR is essential for VMHdm/c^SF-1^ neuronal-induced skeletal muscle *Pgc-1α* expression (Figure 3). β2AdR is expressed throughout the body[18], and even within the skeletal muscle, β2AdR is expressed in a variety of cell types including skeletal muscle cells, smooth muscle cells (blood vessels), and endothelial cells[42; 56]. In addition, recent studies highlight the key role of β2AdR at the neuromuscular junctions (NMJs) to regulate acetylcholine and acetylcholine receptors[57; 58]. As we used global β2 AdR KO mice, we can not formally exclude the possibility that β2AdR in the non-skeletal muscle organs are actually essential. Further studies using tissue specific β2AdR manipulation (e.g., deletion of β2AdR in specifically skeletal or smooth muscle cells) are necessary to decipher the precise targets of VMHdm/c^SF-1^ neurons and the sympathoadrenal axis.

The terminal activation of VMHdm/c^SF-1^ neurons in the PVH/AH and PAG significantly increases blood epinephrine and skeletal muscle *Pgc-1α-2 and -3* mRNA (Figure 4). Because the PVH and PAG directly project to the sympathoadrenal preganglionic neurons in the intermediolateral nucleus of the spinal cord[59–62], it is predicted that activation of these areas can have more profound effects on the sympathoadrenal activity. By comparing the results of the PVH/AH and AH stimulation (Figure 4), it is likely that VMHdm/c^SF-1^ neurons projecting to the PVH substantially contribute to the regulation of the SNS activity than that to the AH. Because ADX completely diminishes augmented blood glucose by VMHdm/c^SF-1^ neuronal stimulation (Figure 2), epinephrine release by the SNS activation is likely required for the augmented blood glucose levels. Interestingly, the terminal activation in all sites we investigated significantly increased blood glucose (Figure 4). Taken together, blood glucose data (Figure 4) indicate that all sites we investigated can activate the sympathoadrenal activity, although the degree of the sympathoadrenal activation may vary. In fact, the mean value of blood epinephrine, skeletal muscle *Pgc-1* mRNA levels in all the sites are higher than control group (Figure 4), suggesting that all sites we investigated contribute to VMHdm/c^SF-1^-neuronal regulation of the sympathoadrenal activity at some extent of degree. Collectively, these data suggest that the downstream sites of VMHdm/c^SF-1^ neurons have redundant functions regarding the augmented sympathoadrenal activity.

The Fos expression data (Figure 5) indicate that VMHdm/c^SF-1^ neuronal axon projections are collaterals rather than one-to-one projections[63] as terminal stimulations can induce back-propagated activation in the proximal projected sites of VMHdm/c^SF-1^ neuronal (Figure 5). A previous study using the retrograde tracing method supports this notion as they found that most of VMHdm/c^SF-1^ neurons project to the AH also collaterally send the axon to the PAG[23]. The VMH regulate essential physiological function for survival and high-energy demand situations including counterregulatory actions to hypoglycemia[3], defensive behavior[23; 40], and exercise[5; 64]. These survival functions have to be executed coordinately at the whole-body level. Considering that collateral VMHdm/c^SF-1^ neuronal circuits are functionally redundant, we propose that similar to monoaminergic neurons[65], VMHdm/c^SF-1^ neurons play a “broadcast” role in the regulation of physiological functions at the whole-body level during emergency or high-energy demand situations. Further studies will be necessary to delineate the degree of contributions of each VMHdm/c^SF-1^ neuronal downstream node to the regulation of metabolism.

## 4.2 Limitation of this study

As many previous studies have noted, optogenetic stimulation induces firing patterns that are dissimilar to endogenous firing patterns in many neurons[66]. Therefore, we can not exclude the possibility that our data demonstrate the maximum capability of VMHdm/c^SF-1^ neurons on the regulation of skeletal muscle rather physiological roles of VMHdm/c^SF-1^ neurons. We also used a fixed optogenetic configuration throughout the terminal activations, despite the fact that each site we investigated has a different density of VMHdm/c^SF-1^ neuronal axon and terminals[21]. It is virtually possible that different terminal sites require different firing pattern to execute their function properly. Future studies to use fine-tuning configuration are necessary to test this possibility.

As we mentioned above, the inherent limitations of the ADX studies have to be considered. Although we supplied the corticosterone to maintain its physiological levels, the surgical removal of the adrenal gland can compromise many physiological functions directly and indirectly. For instance, we observed that the basal levels of skeletal muscle *Pgc-1α* were decreased in ADX mice (56%, 51%, 96%, 95%, and 72% mean reductions in *Pgc-1α total, Pgc-1α-1, Pgc-1α-2, Pgc-1α-3, and Pgc-1α-4* respectively compared to sham control, in Figure 2G). To exclude the possibility that ADX affects the sensitivity of adrenergic receptors, we investigated whether β2AdR agonist can induce skeletal muscle *Pgc-1α* in ADX mice. β2AdR agonist significantly induced skeletal muscle *Pgc-1α*, and there were no significant differences between sham and ADX mice (SFigure 6). Thus, ADX unlikely affects the sensitivity of adrenergic receptors in skeletal muscle in our experimental design. Nonetheless, we have to interpret ADX studies with careful consideration, and future experiments using sophisticated techniques (e.g., genetic-ablation of epinephrine only from the adrenal medulla) will be warranted to further confirm the role of epinephrine releases from the adrenal medulla in the regulation of skeletal muscle physiology.

Fos protein expression is used as the readout of neuronal activity. However, there are limitations we have to consider. First, we can not distinguish whether Fos activation is direct consequences of VMHdm/c^SF-1^ neuronal activation or indirect consequences of physiological changes induced by VMHdm/c^SF-1^ neuronal activation. For instance, it is well appreciated that stressors can increase Fos expressions. VMHdm/c^SF-1^ neuronal activation clearly induce fight or flight type of endocrine responses (e.g., increased epinephrine, corticosterone, glucagon, and glucose in the blood), that potentially can increase Fos expression indirectly.

## 4.3 Conclusion

The CNS-skeletal muscle interactions are important to maintain metabolic homeostasis. The skeletal muscle plays a critical role in the regulation of metabolic homeostasis as it substantially contributes to basal energy expenditure and glucose disposal after meal[13; 67]. A large body of studies has built the foundation of neuroanatomy regarding metabolic homeostasis[62]. In the last decade, optogenetic tools have revealed detailed functional neurocircuits regulating metabolism including food intake and glucose metabolism[1; 68; 69]. This study using optogenetics demonstrates that VMHdm/c^SF-1^ neurons regulate skeletal muscle PGC1-α via epinephrine released from the adrenal gland coupled with β2AdR. In addition, our data suggest that VMHdm/c^SF-1^ neuronal circuits regulating the sympathoadrenal activity are functionally redundant, yet varied contributions from each downstream node to the regulation of the SNS likely exist (SFigure 7). Our study advances the understanding of brain-skeletal muscle communications and implies the significant contributions of VMHdm/c^SF-1^ neurons→sympathoadrenal axis to beneficial effects of exercise on skeletal muscle.

## 5. Acknowledgement

We thank Glenn Toney at UT Health San Antonio (UTHSA) for valuable advice, Nancy Gonzalez and Harun A. Khan (UTHSA), Bandy Chen, Safia Baset, and Jasmine Dushime (UTSW) for the technical assistant, Juri C. Fujikawa for providing animal illustrations, and Steven Wyler, Ryan Reynolds, Luis Leon Mercado, Syann Lee, and Joel. K. Elmquist (UTSW) for valuable comments on the manuscript. This work was supported by the University of Texas System (UT Rising STARs to T.F.), and San Antonio Area Foundation (to T.F.). This manuscript has been released as a pre-print at Biorxiv (https://www.biorxiv.org/content/10.1101/2022.04.01.486756v1).

## 7. Contribution

T.Y. designed, performed and analyzed experiments, and edited the manuscript. M. F., S.F., A.H., Z.S., J.L., A.S.T., and A.K.S., performed experiments. T.F. designed, performed, supervised, and analyzed experiments, and wrote and finalized the manuscript.

---

**Supplemental Figure 1, related to Figure 1.**
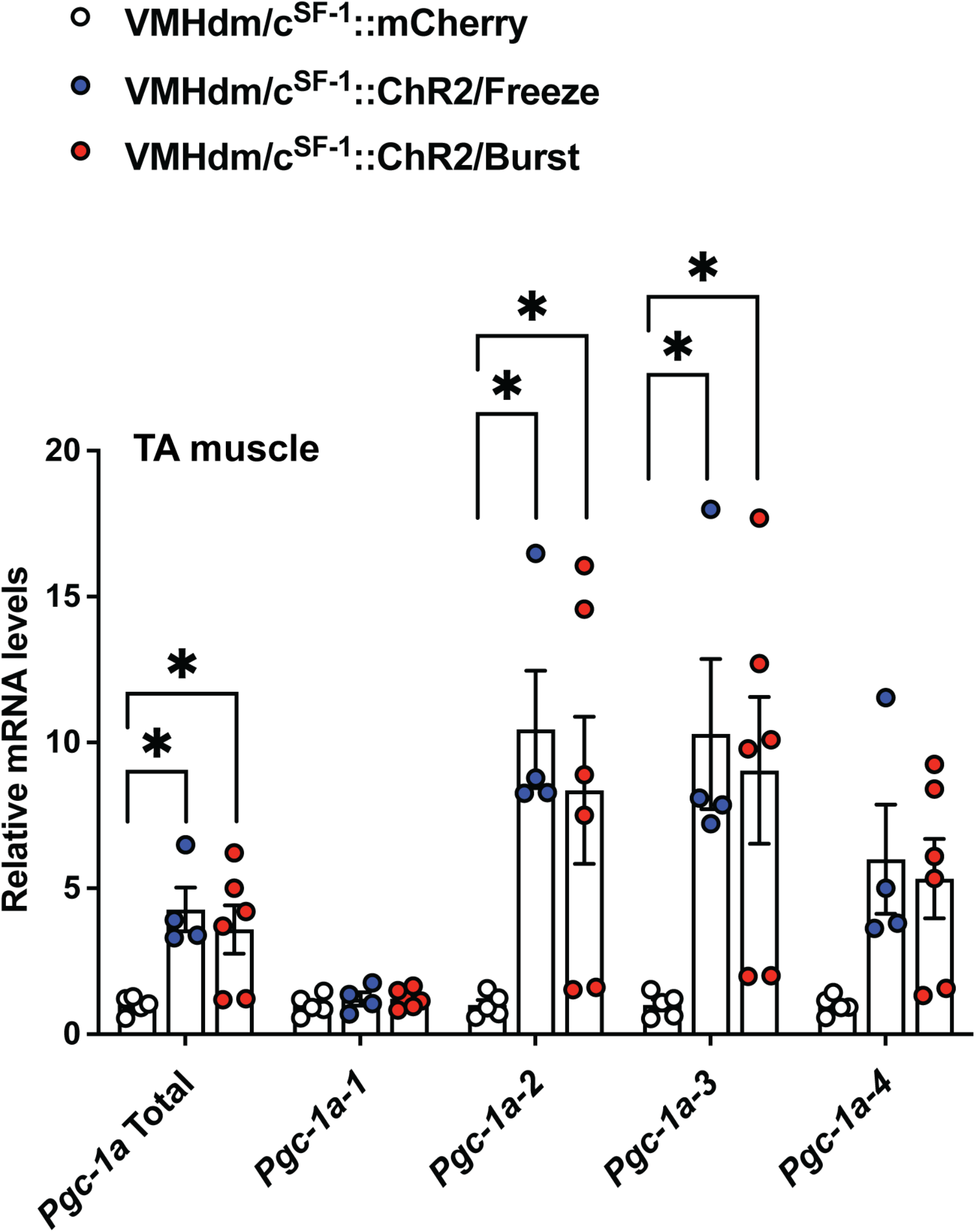
mRNA expression levels of *Pgc1-α* isoform in skeletal muscle of VMHdm/c^SF-1^::ChR2 mice exhibiting freeze or burst behavior after optogenetic stimulation. The stimulation configuration was the same as described in Figure 1. Freeze group was composed of mice that only exhibited freeze behavior. Burst group was composed of mice exhibited burst activity with or without other behavioral changes such as freeze behavior. Values are mean ± S.E.M., *** p < 0.001, * p < 0.05.

**Supplemental Figure 2, related to Figure 5.**
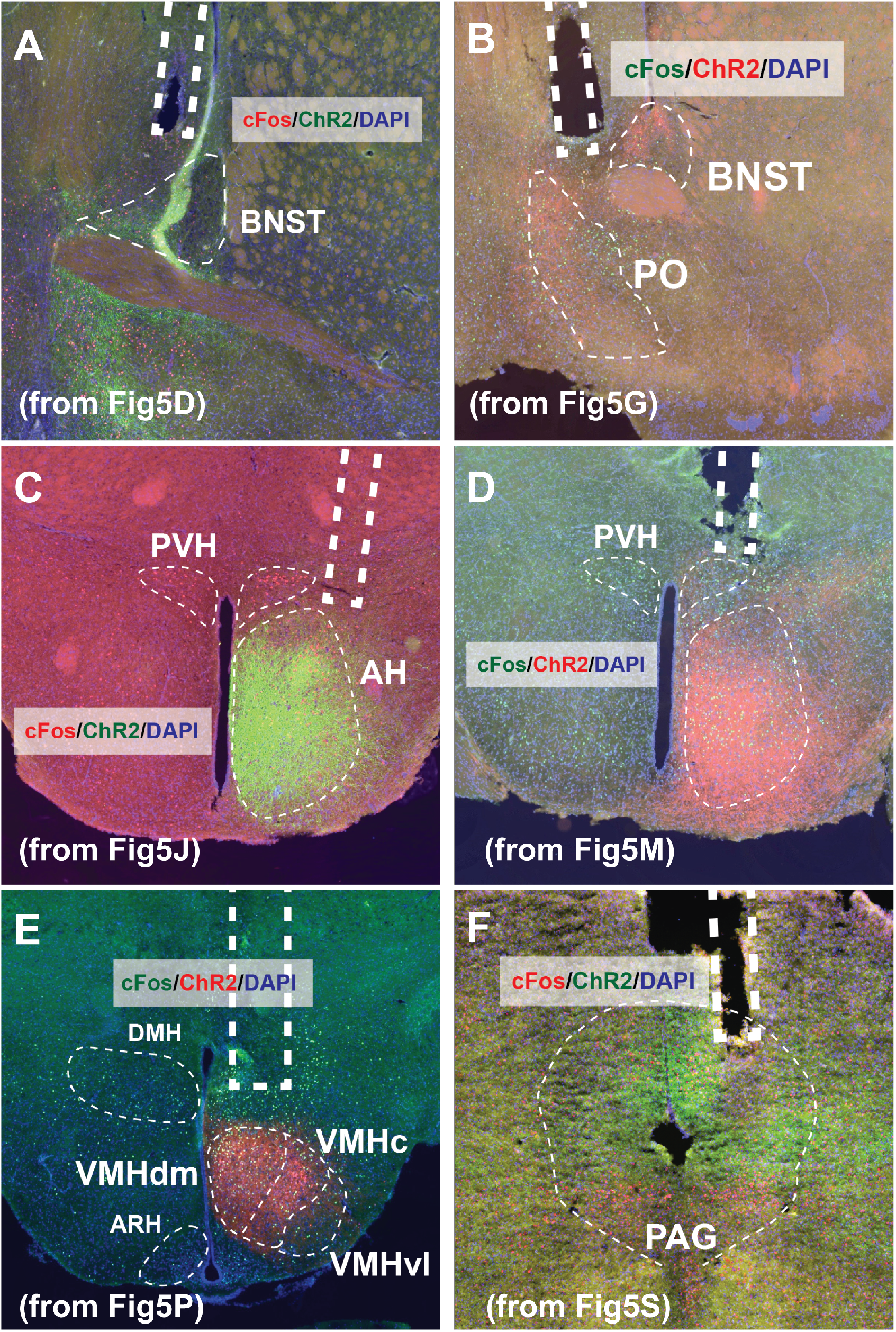
The high-resolution brain maps of optic fiber insertion sites of (**A**), aBNST, (**B**) POA (**C**), AH (**D**), PVH/AH, (**E**), VMH, and (**F**) PAG.

**Supplemental Figure 3, related to Figure 5.**
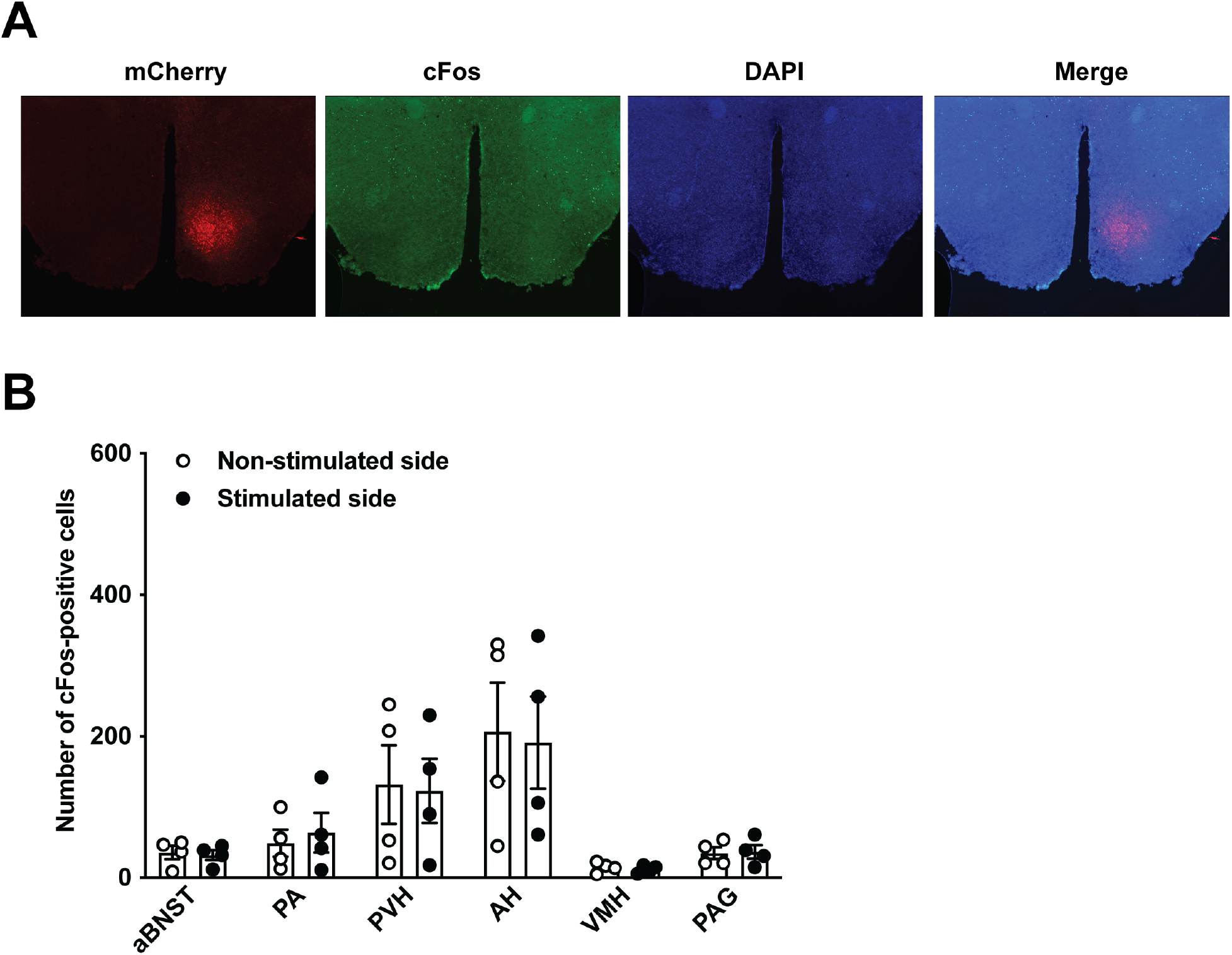
(**A**) Representative figure of the VMH, and (**B**) the number of Fos expression cells at the aBNST, POA, AH, PVH/AH, and PAG of VMHdm/c^SF-1^::mCherry mice after the soma stimulation of VMH.

**Supplemental Figure 4, related to Figure 5.**
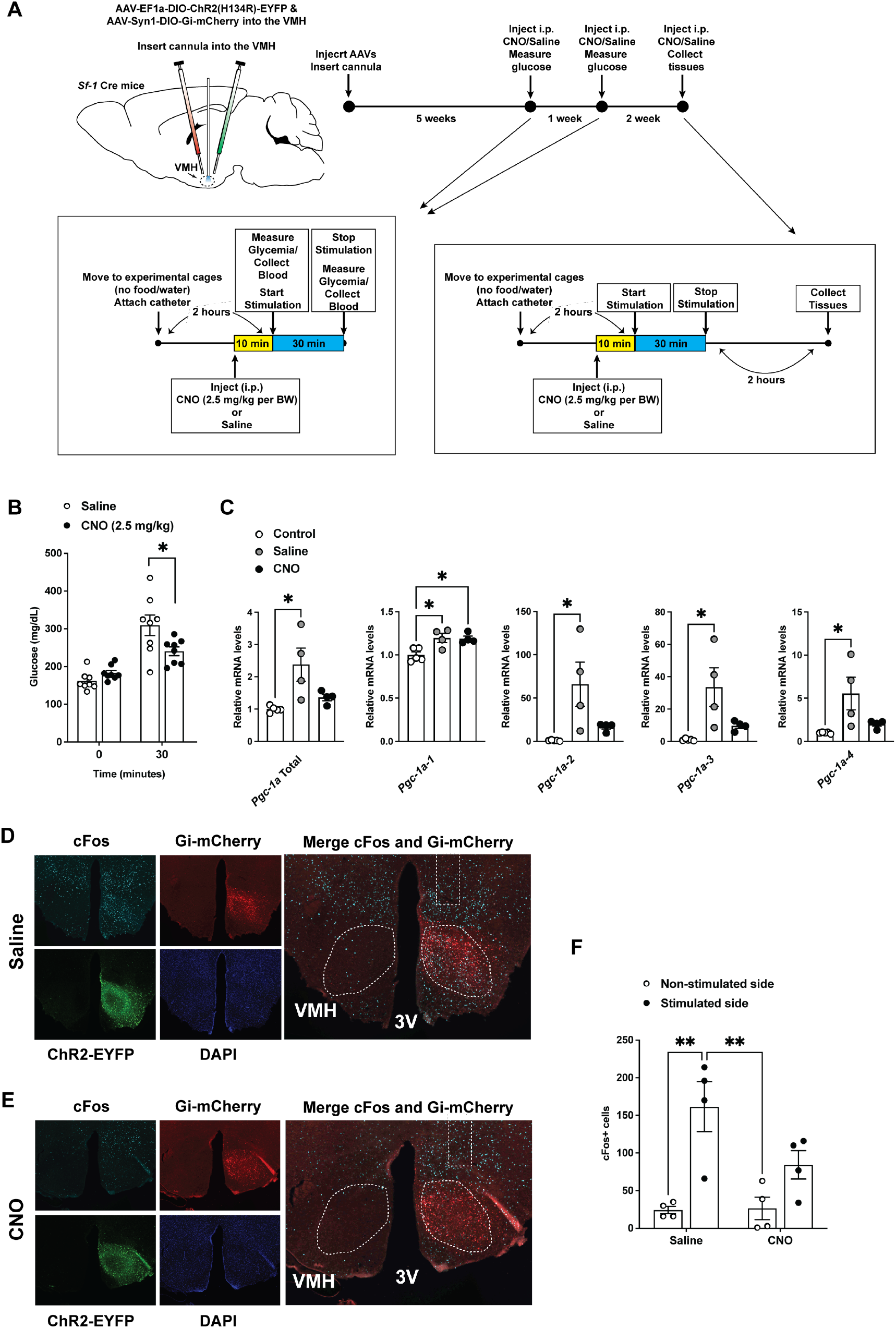
(**A**) Schematic of experimental design. (**B**) Blood glucose levels of Before (0 minutes) and 30 minutes after the VMH stimulation VMHdm/c^SF-1^::ChR2::Gi-DREADD mice. CNO (2.5 mg/kg) was i.p. injected 10 minutes before after the VMH stimulation. Saline was used as control for CNO. (**C**) mRNA expression levels of *Pgc1 isoforms* in skeletal muscle of VMHdm/c^SF-1^::ChR2::Gi-DREADD mice after the terminal stimulation while CNO (2.5 mg/kg) was i.p. injected. Control mice were VMHdm/c^SF-1^::EYFP::Gi-DREADD mice with the VMH stimulation. Representative figures of Fos expression the VMH in VMHdm/c^SF-1^::ChR2::Gi-DREADD mice after the terminal stimulation while (**D**) saline or (**E**) CNO (2.5 mg/kg) was i.p. injected. Magenta, Red, Green, and Blue represents Fos expression, Gi-DREADD, ChR2, and DAPI, respectively. (**F**) Fos number in the VMH in VMHdm/c^SF-1^::ChR2::Gi-DREADD mice after the terminal stimulation while saline or CNO (2.5 mg/kg) was i.p. injected. The stimulation setting was 5 ms duration, 10 Hz, 2 seconds activation and 2 seconds rest cycle, 30 minutes. Values are mean ± S.E.M. ** p <0.01, * p < 0.05.

**Supplemental Figure 5, related to Figure 5.**
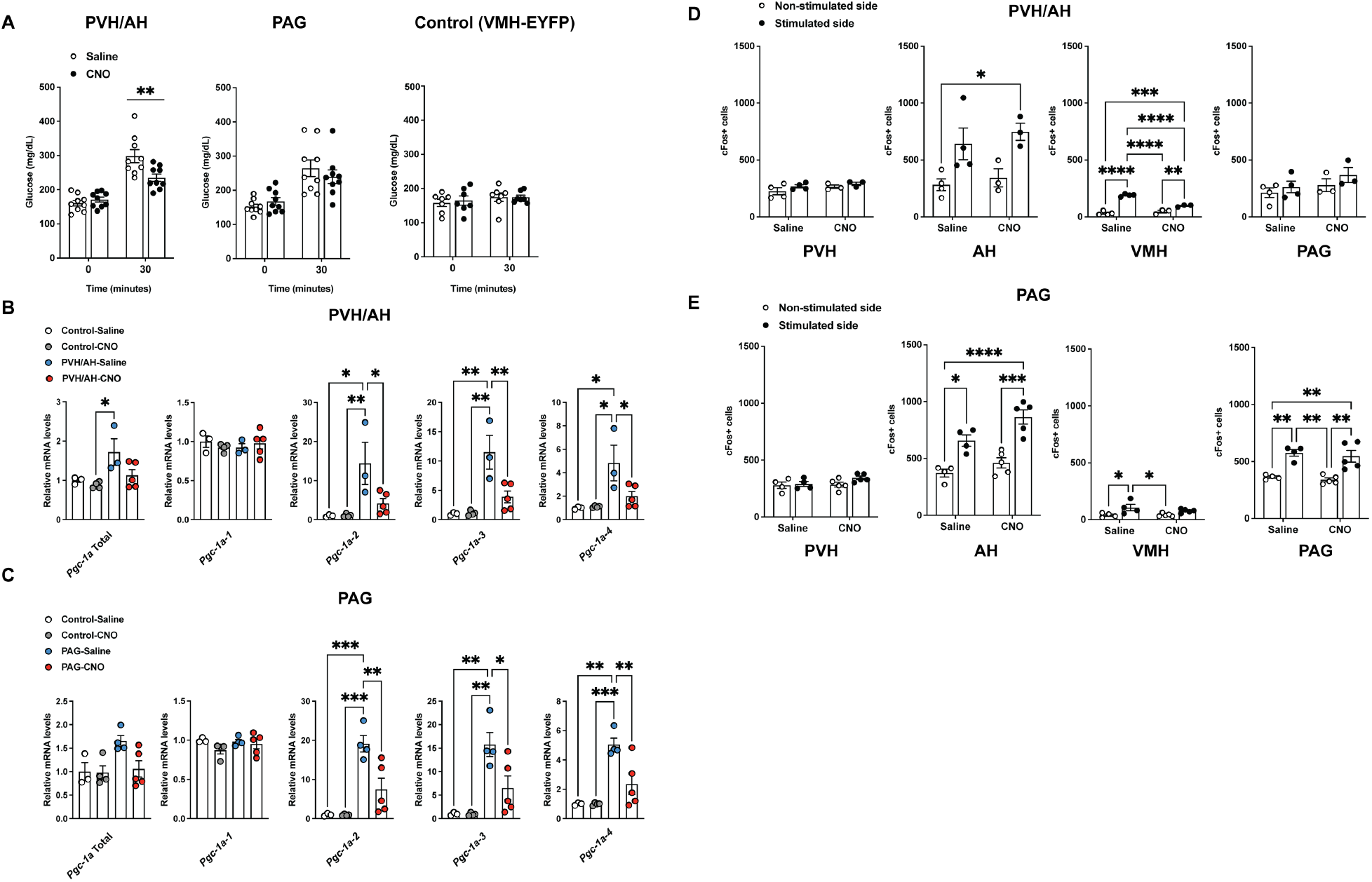
(**A**) Blood glucose levels before (0 minutes) and 30 minutes after the terminal stimulation of VMHdm/c^SF-1^ neurons at the PVH/AH or PAG in VMHdm/c^SF-1^::ChR2::Gi-DREADD mice. CNO (2.5 mg/kg) was i.p. injected 10 minutes before after the terminal stimulation. Saline was used as control for CNO. Control mice for Gi-DREADD were administered rAAV5-EF1α-DIO-EYFP with rAAV8-hSyn-DIO-hM4D(Gi)-mCherry into the VMH of *Sf-1*-BAC-Cre mice. mRNA expression levels of *Pgc1 isoforms* in skeletal muscle of VMHdm/c^SF-1^::ChR2::Gi-DREADD mice after the terminal stimulation of VMHdm/c^SF-1^ neurons at the (**B**) PVH/AH or (**C**) PAG while CNO (2.5 mg/kg) was i.p. injected. Control mice were VMHdm/c^SF-1^::EYFP::Gi-DREADD mice with the VMH stimulation. Fos number in the PVH, AH, VMH, and PAG in VMHdm/c^SF-1^::ChR2::Gi-DREADD mice after the terminal stimulation at the (**D**) PVH/AH or (**E**) PAG while saline or CNO (2.5 mg/kg) was i.p. injected. The stimulation setting was 5 ms duration, 10 Hz (PVH/AH) or 20 Hz (PAG), 2 seconds activation and 2 seconds rest cycle, 30 minutes. Values are mean ± S.E.M. **** p < 0.0001, *** p < 0.001, ** p <0.01, * p < 0.05.

**Supplemental Figure 6, related to Figure 2.**
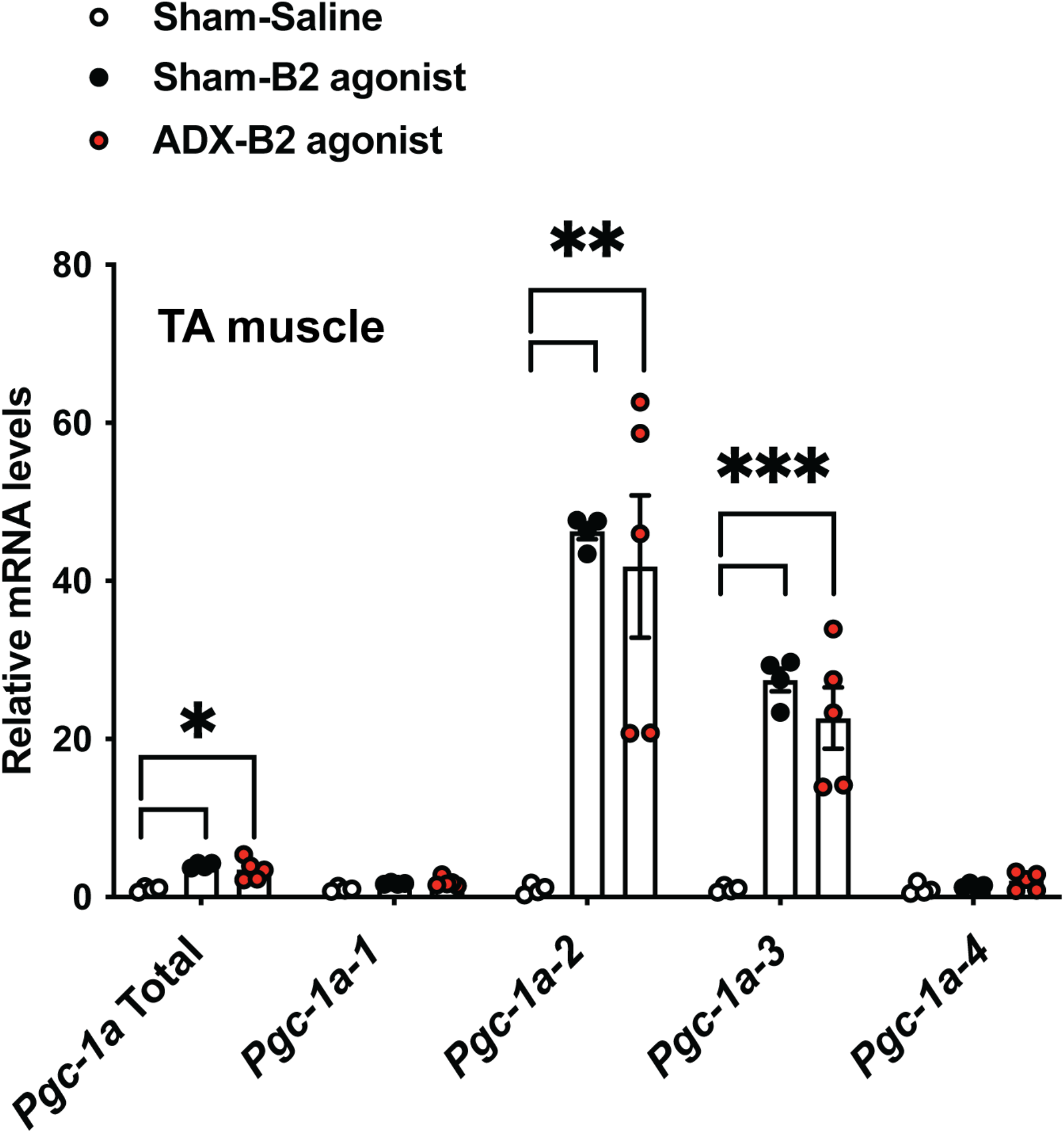
mRNA expression levels of *Pgc1-α* isoform in skeletal muscle of ADX mice after β2AdR agonist (clenbuterol 1 mg per kg bodyweight, dissolved in sterile saline solution). Values are mean ± S.E.M. *** p < 0.001, ** p <0.01, * p < 0.05.

**Supplemental Figure 7.**
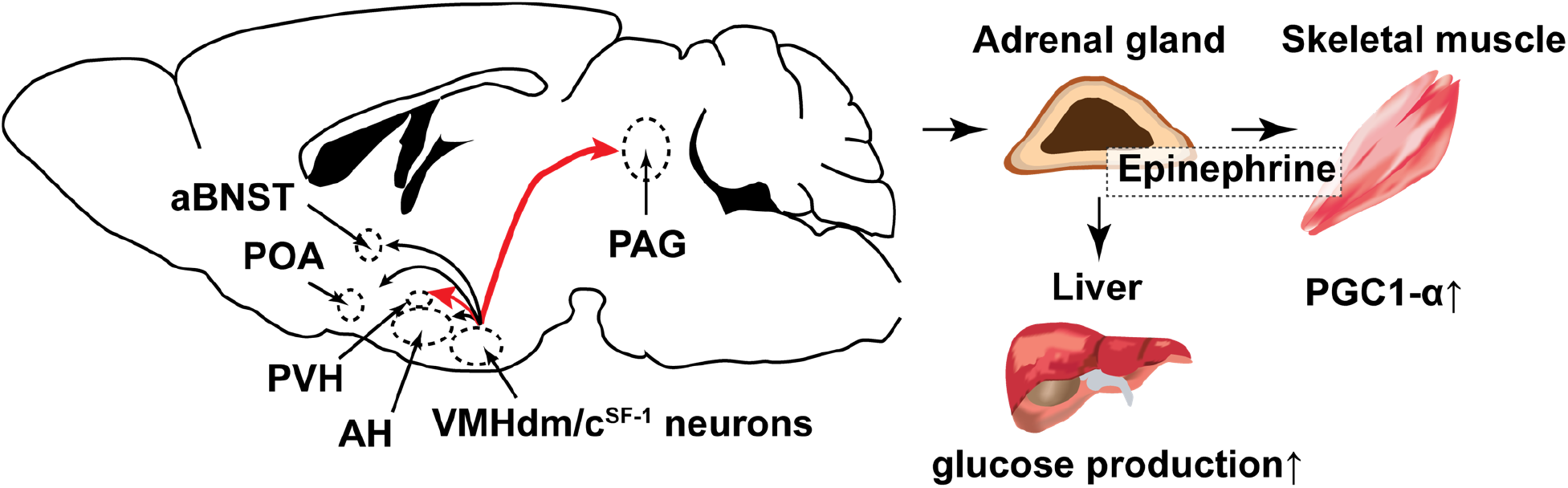
Summary figure depicting pathways by which VMHdm/c^SF-1^ neurons regulate skeletal muscle *Pgc1-α* expression and blood glucose levels via the sympathoadrenal gland.

